# A visual paired associate learning (vPAL) paradigm to study memory consolidation during sleep

**DOI:** 10.1101/2023.03.28.534494

**Authors:** J.F. Schmidig, M. Geva-Sagiv, R. Falach, S. Yakim, Y. Gat, O. Sharon, I. Fried, Y. Nir

## Abstract

The hippocampus helps transform an experience into an enduring memory by associating its multiple aspects. Sleep improves the consolidation of the newly formed associations, leading to stable long-term memory. Most research on human declarative memory and its consolidation during sleep uses word-pair associations requiring exhaustive learning. Here we present the visual paired association learning (vPAL) paradigm, in which participants learn new associations between images of celebrities and animals. vPAL associations are based on a one-shot exposure that resembles learning in natural conditions. We tested if vPAL can reveal a role for sleep in memory consolidation by assessing the specificity of memory recognition, and the cued recall performance, before and after sleep. We found that a daytime nap improved the stability of recognition memory and discrimination abilities compared to identical intervals of wakefulness. By contrast, cued recall of associations did not exhibit significant sleep-dependent effects. High-density EEG during naps further revealed an association between sleep spindle density and stability of recognition memory. Thus, the vPAL paradigm opens new avenues for future research on sleep and memory consolidation across ages and heterogeneous populations in health and disease.

## Introduction

New memories are initially weak, but such labile memories are further strengthened and integrated into pre-existing long-term memory through consolidation. Accumulating evidence established that sleep is critical for memory consolidation (Datta & MacLean, 2007; Diekelmann & Born, 2010; Dudai et al., 2015; Maquet, 2001; Tononi & Cirelli, 2014; Walker & Stickgold, 2004). Over time, it has been repeatedly confirmed that sleep supports memory consolidation (Diekelmann & Born, 2010) and that sleep-related benefits in learning are not simply due to lower fatigue, circadian time, or reduced interference. Numerous studies confirmed the beneficial effect of sleep memory in various tasks including motor, procedural, perceptual, and emotional memories (Fischer et al., 2002; Karni et al., 1994; S. Mednick et al., 2003; Payne et al., 2008; Robertson et al., 2004; Wagner et al., 2001; Walker et al., 2005). Significant benefits of sleep on memory are observed after an 8-hour night of sleep, but also after shorter naps of 1– 2 hours (Korman et al., 2007; S. Mednick et al., 2003; Nishida et al., 2009; Tucker et al., 2006).

Among the many different memory systems examined, the consolidation of declarative memory has received much attention in the past years (for a review see Cordi & Rasch, 2021; Marshall & Born, 2007). The first demonstration that sleep benefits memory was obtained in 1924 using the paired associate learning (PAL) paradigm in humans (Jenkins & Dallenbach, 1924). Since then, it has been repeatedly demonstrated that the transformation of these initially weakly bound associations into long-lasting memories benefits from sleep (Barrett & Ekstrand, 1972; Gais et al., 2006; Lahl et al., 2008; Lau et al., 2010; Plihal & Born, 1997; Rasch et al., 2007; Ruch et al., 2012; Studte et al., 2015; Tucker et al., 2006).

Sleep not only contributes to memory consolidation, but certain sleep events are specifically associated with benefits for long-term memory. During a participant’s slumber, specific electrophysiological aspects of NREM sleep, e.g. sleep spindles, are crucial role for memory consolidation (Gais et al., 2002; Marshall et al., 2006; Mednick et al., 2013; Mölle et al., 2011; Schabus et al., 2004). Beyond correlation, a pharmacological manipulation that increases sleep spindles improves verbal memory performance, whereas diminishing spindles impaires memory performance (Mednick et al., 2013)

Nearly all previous studies on human sleep and hippocampus-dependent declarative memory used a word-pair learning task where two nouns (e.g. flowerpot and airplane) are paired, and memory retrieval is tested after sleep either by recognition and/or by free/cued recall. This approach follows a tradition of verbal learning and memory (Hall, 1971) but, despite its wide utility, relying solely on this specific semantic-oriented task also has disadvantages. First, it has been argued that the failure to replicate some key findings on sleep and memory may stem from the fact that effects depend on the specific task used (Bailes et al., 2020; Cordi & Rasch, 2021; Pöhlchen et al., 2021). Furthermore, it has been shown that laboratory-based and more natural autobiographical retrieval substantially differ in their neural substrates (McDermott et al., 2009). Thus, expanding experiments to use other closer-to-real-life tasks would potentially improve robustness in this research domain. Second, word-lists are often difficult to memorize, thereby evoking weaker memory performance. Accordingly, learning such lists typically requires repeated encoding and extensive rehearsal before reaching an adequate performance level. Such rehearsal inevitably involves factors such as motivation that go beyond memory per-se. Third, using word pairs to study the relation between sleep and memory consolidation may be suboptimal for research in elderly and clinical populations (e.g. dementia) given that learning word-pair lists depends on reading speed and attention span that are often impaired in such individuals.

To go beyond some of these limitations, it would be beneficial to expand the toolset for research on sleep and declarative memory consolidation beyond the classical word pair task. A first step are face-name/word association paradigms, which have exchanged one word of the association pair with a picture (Ellenbogen et al., 2007; Niknazar et al., 2022; Ruch et al., 2012). However, language remained a prerequisite to learn new associations. Hence, we developed the visual paired associate learning (vPAL) paradigm: a short (∼15-minutes), image-based memory paradigm with high ecological validity. Our objective was to design a paradigm that encompasses more aspects of natural memory formation relative to the frequently used word lists. To this end, asked participants to form new associations between image pairs of celebrity faces and animals. Mimicking everyday-like experiences, this task is based on one-shot learning. Moreover, by presenting familiar and engaging images, the vPAL paradigm is well suited for clinical populations because it is time-efficient (no rehearsal), less reliant on reading, and easy to comprehend. We set out to test how sleep affects memory consolidation in this novel vPAL paradigm by employing a within-subject design in young healthy volunteers. Each participant learned and retrieved associations before and after an interval of either wake or sleep (in two separate sessions), to investigate the effects of sleep beyond those of time elapsed or circadian factors.

## Results

To investigate the possible effects of sleep on memory consolidation with a visual paired associate learning (vPAL) paradigm, healthy young volunteers (n = 28, average age = 27.0 years, range: 22 to 37 years, 39% female) participated in two sessions taking place one week apart (**Figure 1A**, Methods). Each session started with a short (∼15 min) encoding session around noon, where participants learned 25 pairings between photographic images of famous people and animals (contextualized as “pet owners” and their pets). Learning was followed by an ‘immediate retrieval’ test, and an identical ‘delayed retrieval’ test, separated by 2.5 hours during which participants were either asked to nap while monitored with polysomnography and high-density EEG or stayed awake. The order of sleep/wake sessions was counterbalanced, and a different image set was used in each session.

**Figure 1.**
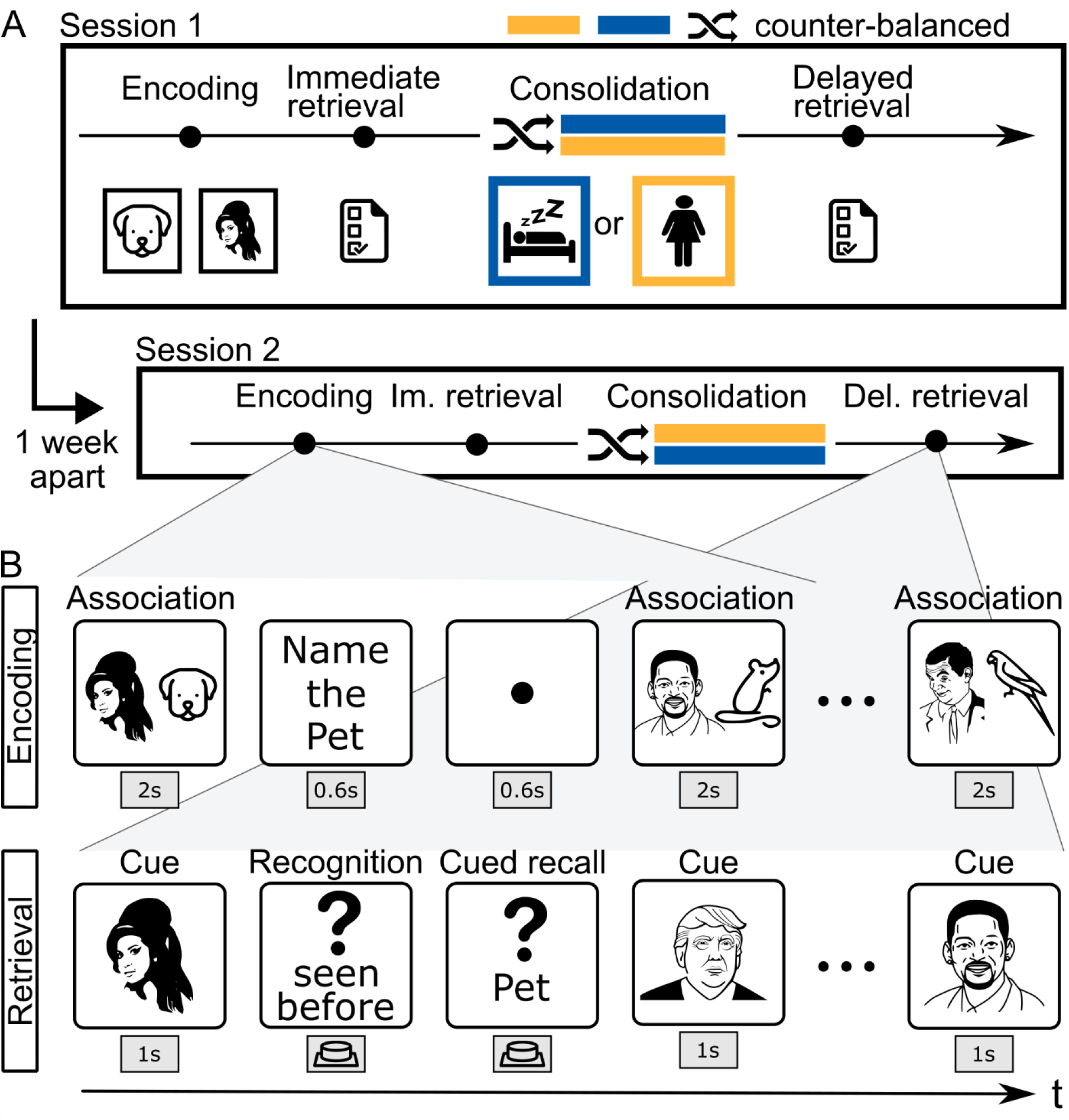
Study design: **(A)** Session design: participants completed two sessions, each including an encoding phase, a consolidation period, and two retrieval tasks (immediate and delayed). The two sessions took place one week apart. We varied the type of consolidation between the sessions, i.e. participants either stayed awake (yellow) or slept (blue) (2.5 hours; order was counterbalanced). **(B)** Task description: encoding and retrieval consisted of the following parts. Encoding phase: associative pairing of celebrity and pet was followed by naming the animal type. Retrieval tasks (immediate and delayed): participants were presented with a celebrity image (cue); then were asked to identify whether the celebrity was earlier paired with a pet or not (recognition) and finally asked to name the specific pet that was paired with the presented celebrity (cued recall).

To verify that participants had adequate sleep, we first performed sleep scoring according to established criteria (AASM, (Iber et al., 2007) using a combination of electroencephalography (EEG), electromyography (EMG), electrooculography (EOG) and video (see methods). Next, two aspects of memory performance were assessed in each retrieval session. First, the specificity of recognition memory (‘Was this celebrity seen before?’) was evaluated via (yes/no) responses to previously learned images and a set of lures (the same set of lures was used in immediate and delayed retrieval). We quantified ***recognition discrimination*** as the difference between hit-rate and false-detection rate to provide a measure of item memory (Methods). Second, we evaluated the rate of ***correctly recalled associations*** (‘What was the pet of this celebrity?’) as the percentage of correctly named pets. Performance on both measures was assessed without providing feedback to any of the responses four times for each participant (2 time points * 2 sessions): in the late morning (immediate retrieval, M = mean (standard deviation): M = 13:15 (1:09)) and the afternoon (delayed retrieval, M = 16:13 (1:16)), separately in sessions with a nap opportunity or in sessions with wakeful rest.

### vPAL memory performance declines over time

Participants successfully learned the celebrity-pet associations. In the immediate retrieval session, they recognized 88.7 ± 13.5% (mean ± SD) of previously presented celebrities (significantly more than chance, p < 0.0001), and correctly associating some of the corresponding pets (38.4 ± 21.3%, p < 0.0001). We compared memory performance in the immediate and delayed retrieval and found that, as expected (Ebbinghaus, 1913), memory performance declined over time in all experimental conditions (**Fig. 2**).

**Figure 2.**
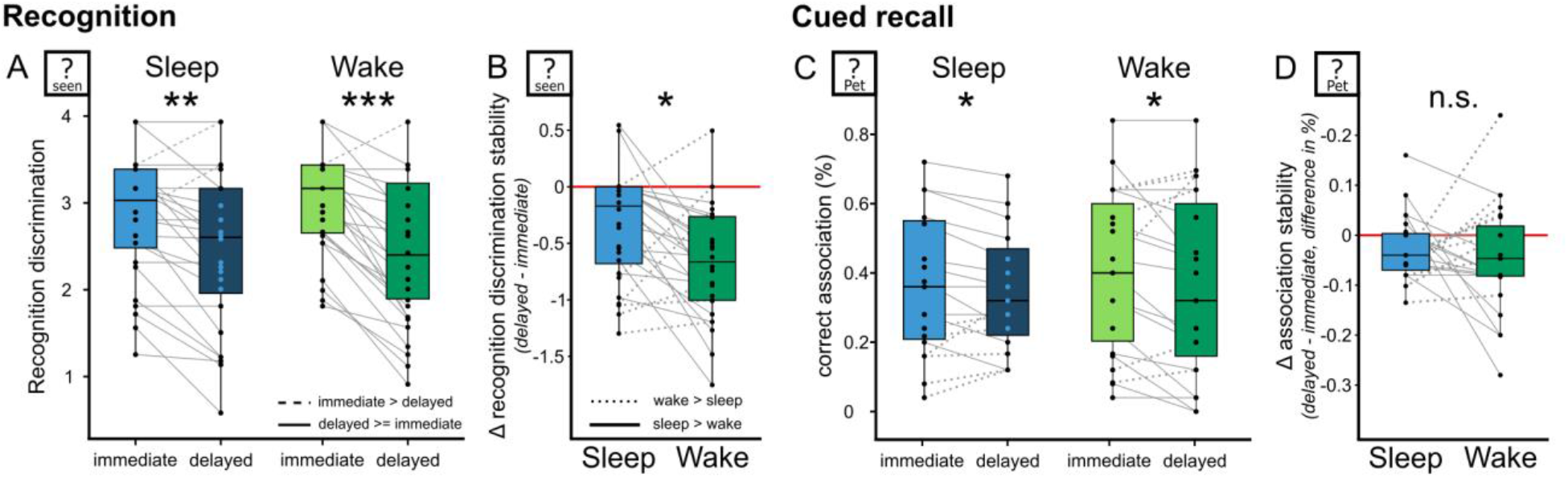
Sleep benefits memory recognition performance: Difference in memory retention for the recognition task (left) and the cued recall (right), compared between immediate and delayed retrieval, as well as between consolidation type (wake vs sleep). **(A)** Recognition discrimination before (light color) and after (dark color) in the sleep (left) or wake condition (right). Boxplots depict memory discrimination (y-axis) and retrieval session (x-axis). The horizontal black line marks the median. The box contains the 25^th^ to 75^th^ percentile the vertical black line marks the 5^th^ and 95^th^ percentile limits. Dots depict individual subjects (n = 28) and their performance in the two separate sessions. Full (and dotted) lines indicate whether memory performance of the same participant was higher (or lower) in the immediate compared to the delayed retrieval session. **(B)** Recognition discrimination stability (delayed-immediate difference in performance, y-axis) in wake (green) or sleep (blue) sessions. Here full (and dotted) lines indicate whether a participant’s memory preservation was higher (or lower) after sleep than after wake rest. Red line indicates stable memory (no memory decline between immediate and delayed recall). **(C)** Same as **A** and **(D)** same as **B** for successful association in cued recall (n = 23). * = p < 0.05, ** = p < 0.01, *** = p < 0.001, n.s. = not significant.

Recognition discrimination declined between the first and second retrieval tests conducted 2.5 hours apart. The recognition discrimination declined on average by 16.5 ± 18.6% over time (**Fig. 2A**, Difference_delayed-immediate_ = −0.482 ±-0.544, Wilcoxon signed-rank test V = 79, p < 0.0001, Cohen’s d = 0.89, recognition memory: M _immediate_ = 2.93 (0.66), M _delayed_ = 2.44 (0.9)). This robust decline was also observed when we analyzed the data for the sleep and the wake rest group independently (for both conditions V > 15; p < 0.002). The amount of correctly recalled association also declined over time. Participants remembered significantly more celebrity-associated pets in the immediate retrieval session compared with the delayed retrieval session (**Fig. 2C**, Wilcoxon signed-rank test V = 187.5, p < 0.005, Cohen’s d = 0.34, recall memory: M _immediate_ = 2.93 (0.66), M _delayed_ = 2.44 (0.9)). A robust decline in recalled associations was also observed when examined separately either for wake or sleep conditions (V > 40; p < 0.05). These results demonstrate that memory performance declined over time.

### vPAL memory recognition is better conserved after sleep than after wakefulness

To assess whether sleep benefited memory consolidation, we compared whether memory performance was better maintained after sleep than after an equally long interval of wakefulness. We found that recognition discrimination was significantly better conserved after sleep (**Figure 2B**, t(27) = 2.51, p _paired_ = 0.018, Cohen’s D = 0.47, Δ recognition discrimination stability: M _sleep_ = −0.34 (0.48), M _wake rest_ = −0.63 (0.57)). This effect size is at the upper end of what has been described for sleep-mediated memory effects (Cordi & Rasch, 2021). By contrast to recognition memory, the cued recall of the celebrity-pet associations did not show significant better stability of successful associations after sleep (**Figure 2D**, Δ association stability: t(22) =0.64, p = 0.526, Cohen’s D = 0.13, M _sleep_ = −0.03 (0.07), M _wake rest_ = −0.04 (0.11)). Thus, measures of recognition memory are better conserved after sleep, while cued associations did not exhibit significant sleep-associated improvements.

### vPAL memory recognition correlates with sleep spindle density

We proceeded to go beyond considering sleep in an ‘all or nothing’ manner. First, we examined how variability in sleep architecture and sleep spindle activity correlated with individual changes in memory performance. **Figure 3A** shows an example hypnogram (time-course of sleep stages) throughout the 150min (2.5h) period when sleep was allowed. On average, subjects had a total sleep time (TST) of 118.4 ± 20.2 minutes (mean ± SD), representing sleep efficiency (SE, sleep time per time in bed) of 78.3 ± 12.7 % with a sleep latency of 8.2 ± 8.5 minutes. Average NREM sleep, REM sleep, and wake after sleep onset (WASO) constituted 101.1 ± 19.3, 17.2 ± 12.7, and 17.8 ± 13.6 minutes of sleep time, respectively. Within NREM sleep, N1, N2, and N3 constituted 23.0 ± 20.0, 46.7 ± 23.4, and 31.6 ± 17.6 percentage time after sleep onset, respectively (**Figure 3B**). These features represent a typical profile for a daytime nap in healthy young volunteers (S. Mednick et al., 2003). We then correlated sleep architecture features with the effect of sleep on memory recognition (immediate-delayed retrieval in the sleep group). No correlations emerged as statistically significant, but we observed non-significant trends for positive association between N2 duration and memory recognition discrimination (N2 and memory: R= 0.24, p = 0.221) and negative association between intermittent sleep and shallow N1 sleep and memory recognition discrimination (WASO and memory: R = −0.28, p = 0.144; N1 and memory: R = −0.22).

**Figure 3.**
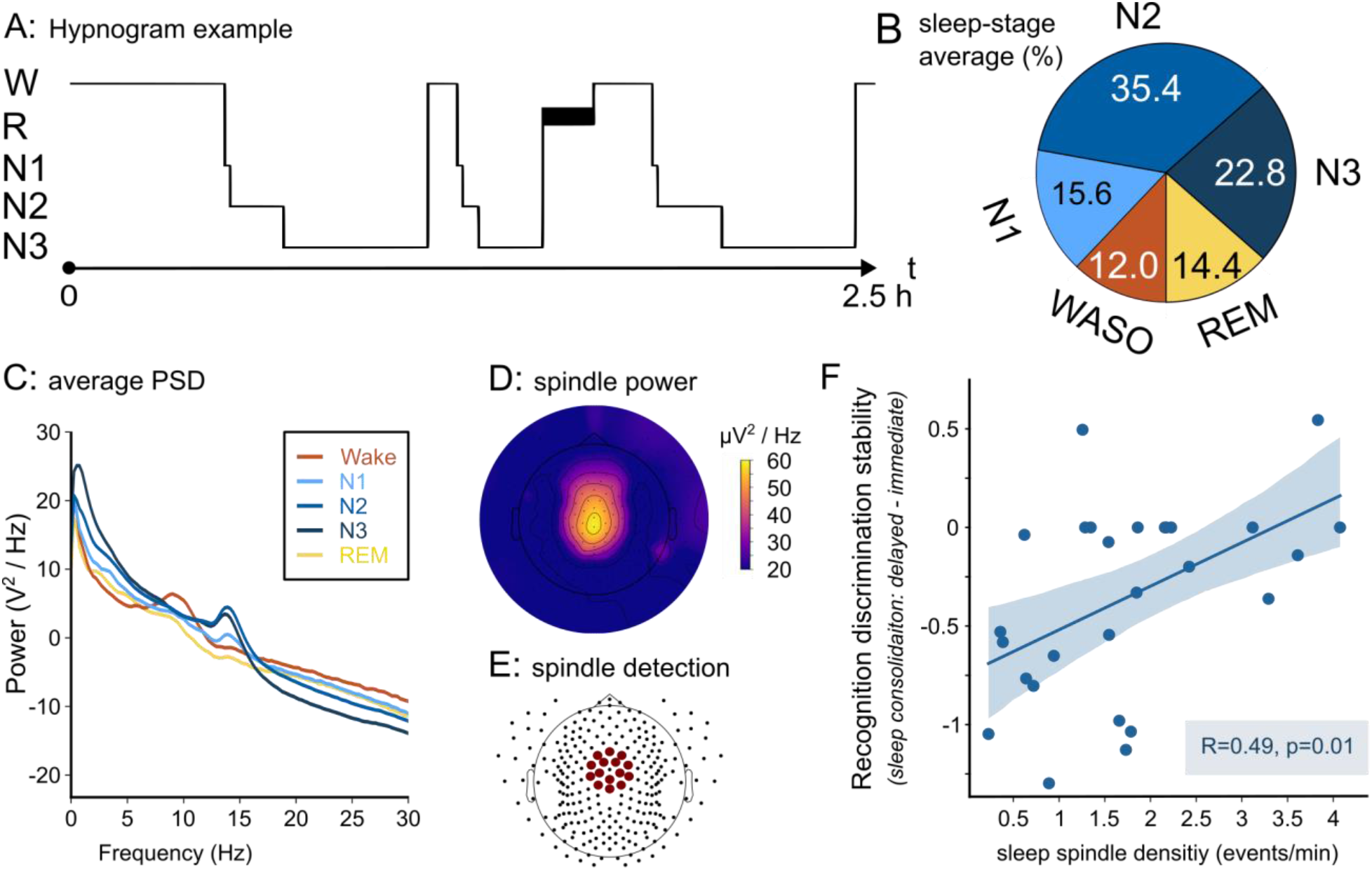
Sleep with high sleep spindle density correlates with better memory recognition discrimination. **(A)** Representative example of a hypnogram (progression of sleep stages on y-axis as a function of time on x-axis) in one participant. W = wake, R = REM sleep, N1-N3 = NREM sleep stages 1-3. **(B)** Pie chart showing average sleep stage distribution across all participants (N=28). WASO = wake after sleep onset. **(C)** Average EEG power spectral density (PSD) per vigilance state for electrode Pz. Note alpha (∼10Hz) and high-frequency activity in wake, slow-wave (<4Hz) and sigma (sleep spindle 10-15Hz) activities in NREM sleep. **D)** Average EEG power in the fast spindle range (13-16Hz), averaged over N2 & N3 sleep of all participants. **(E)** Centro-parietal electrodes used for spindle detection. **(F)** A scatter plot showing recognition discrimination stability after sleep (Δ delayed – immediate recognition rate, y-axis) as a function of sleep spindle density (x-axis). A significantly positive correlation (R _pearson_ = 0.49, p = 0.01) indicates that the vPAL is effective in exposing the association between memory consolidation and sleep spindle activity.

Next, we went beyond sleep architecture to examine in more detail the sleep EEG during nap sessions. EEG power spectrum exhibited expected signatures of each vigilance state including alpha and high-frequency activities in wakefulness, slow-wave activity in N3 sleep, sigma (10-16Hz) activities corresponding to sleep spindles in both N2 and N3 sleep, and diffuse theta activity in REM sleep (**Figure 3C**), as in overnight sleep (Sharon & Nir, 2018). We then tested whether the known association between sleep spindles and successful memory consolidation (Cairney et al., 2018; Diekelmann & Born, 2010; Seibt et al., 2017; Walker & Stickgold, 2004), can be observed with the vPAL paradigm. To this end, we detected sleep spindle events with the YASA algorithm (Vallat & Walker, 2021), focusing on centro-parietal fast (13-16Hz) spindle events occurring in N2/N3 sleep (Methods, **Figure 3D, E**), which are most implicated in hippocampal-dependent memory consolidation (Bar et al., 2020; Petzka et al., 2022; Studte et al., 2015; Zeitlhofer et al., 1997)). We detected 1.72 ± 1.07 events per minute, in agreement with previous studies (e.g. (Andrillon et al., 2011; Nir et al., 2011; Nishida & Walker, 2007; Zeitlhofer et al., 1997). We found a significant positive correlation between spindle density in each individual and the difference of their recognition discrimination befor and after sleep (**Figure 3F**, R = 0.49 p = 0.008). The retention rate before sleep did not correlate with the spindle density (R = 0.014, p = 0.95), hence it is unlikely that our correlation was driven by a general relationship of individuals’ learning and sleeping capabilities. Put simply, the more spindles a participant exhibited per minute; the more their memory was preserved. Altogether, the results show that the vPAL paradigm is effective in revealing the beneficial role of sleep in memory consolidation, and specifically sleep enriched with fast sleep spindles.

## Discussion

We tested if and how sleep affects memory consolidation in a novel visual paired associate learning (vPAL) paradigm. We found that memory decay was less pronounced after sleep, compared to identical intervals of wakefulness, reflecting better memory consolidation. Sleep benefited performance for memory recognition discrimination, but this was not observed for cued recall of associations. Individual recognition discrimination performance was positively correlated with each participant’s sleep spindle density during NREM sleep.

The vPAL holds promise as a new tool that can enrich the experimental toolset for the investigation of sleep and memory consolidation, above and beyond widely used PAL tasks using word pairs (Barrett & Ekstrand, 1972; Ellenbogen et al., 2006; Gais et al., 2006; Lau et al., 2010; Plihal & Born, 1997; Tucker et al., 2006; Werchan & Gómez, 2013). Given that sleep’s effects on memory may depend on specific conditions (Cordi & Rasch, 2021), vPAL could help shed light on how general sleep-mediated memory benefits are - beyond those observed with one specific but widely used task. In this context, our results replicate that sleep benefits memory consolidation in healthy subjects, and those benefits are observed using an image-based PAL task. The vPAL task does not include rehearsal or feedback, to resemble a natural encoding process. Despite the absence of any rehearsal and its short duration, vPAL exhibits an effect size that is higher than most effects reported in studies of human sleep and declarative memory (Cordi & Rasch, 2021). In addition to its effectiveness, the intuitive and engaging nature of vPAL makes it particularly suitable for clinical applications where learning, reading, attention, and rehearsal abilities may be limited. However, some limitations of the study should also be acknowledged. The ecological design leads to a low number of stimuli (25) and we cannot exclude biases due to varying levels of prior celebrity knowledge among the participants. Finally, we focused on a daytime nap which may not fully represent the effects of overnight sleep.

The main result observed here is the reliability with which the vPAL is able to detect the beneficial effect of sleep on memory consolidation. Consolidation during sleep, when compared to wakefulness, improved the ability to differentiate previously presented celebrities versus lure images, an aspect that is often associated with hippocampal function (Quiroga et al., 2005; Suthana et al., 2015). Since we tested celebrities, participants were familiar with the faces prior to the study. By cueing pre-existing, conceptual information, we opted to investigate hippocampus-dependent memory specificity that goes beyond extra-hippocampal familiarity processes (Bird, 2017; Smith et al., 2014).

In addition to investigating memory consolidation during sleep versus wakefulness, our findings establish that vPAL reveals correlation between memory improvement during sleep and sleep spindle rate. This finding joins the growing body of evidence supporting the association between sleep spindles and memory consolidation (Bar et al., 2020; Cairney et al., 2018; Gais et al., 2002; Hennies et al., 2016; Schabus et al., 2004; Studte et al., 2015), and aligns well with the hypothesis that sleep spindles facilitate synaptic plasticity (Seibt et al., 2017; Sejnowski & Destexhe, 2000) by representing an offline reactivation of previously acquired knowledge (Antony et al., 2019; Bergmann et al., 2012; Muehlroth et al., 2019; Petzka et al., 2022; Schönauer et al., 2017). A recent study (Geva-Sagiv et al., 2023) reported vPAL memory benefits in epilepsy patients upon intracranial stimulation coupled to hippocampal slow-wave up-states. Importantly, the degree of memory improvement was highly correlated with sleep spindle activity, in agreement with the current findings in healthy volunteers. Together, our results suggest that the vPAL paradigm is effective in revealing the involvement of sleep spindles in memory consolidation during sleep.

The beneficial role of sleep in the consolidation of declarative memory is believed to be mediated by interactions between hippocampal and neocortical networks (Buzsáki, 1989). While new memories are initially encoded by the help of the hippocampus, they are gradually integrated into long-term memory without compromising pre-existing memories (Frankland & Bontempi, 2005; Káli & Dayan, 2004; McClelland et al., 1995), a process that occurs preferentially during sleep and rest. During NREM sleep, driven by the slow oscillations that originate in the neocortex (Steriade & Timofeev, 2003), newly encoded information in the hippocampus is repeatedly reactivated in conjunction with sharp wave–ripple activity and thalamocortical spindle activity (Buzsáki, 1989; Geva-Sagiv et al., 2023; Nádasdy et al., 1999; Peyrache et al., 2009; Rudoy et al., 2009). Hence the association between spindles and memory improvement found in this study is consistent with the previously reported benefits of spindle events for sleep-related memory formation. Furthermore, they might mark brief moments of memory reactivation, facilitating the transformation of the newly encoded information from hippocampal to neocortical dependence.

One intriguing aspect of the vPAL-results is that sleep benefited memory recognition but was not associated with a significant improvement in cued recall. The absence of a beneficial effect of sleep on the recall of associations has also been observed in a separate cohort of epilepsy patients performing this task before and after overnight sleep (Geva-Sagiv et al., 2023), as well as in a different face-name associations task (Backhaus & Junghanns, 2006). However, we cannot exclude the possibility that sleep also affects recalling celebrity–pet associations. It could be that, with our specific experimental design, the cued recall task was not sufficiently sensitive to detect differences evoked by sleep, but that such differences do exist. Along this line, the initial retrieval rate immediately (minutes) after learning was around one-third of the associations (Figure 2C). With this low success rate, sporadic lapses of attention or other noise sources could mask the potential influence of sleep on memory consolidation.

An alternative possibility is that the results represent a genuine difference in how sleep affects the consolidation of visual recognition memory vs. associations. In this case, the results may be at odds with other (i.e. verbal rather than visual) PAL studies reporting greater benefits of sleep for recall than for recognition (Newbury & Monaghan, 2019). If true, such a discrepancy would pose an open question and, at this point, we can only speculate about how and why these two aspects of memory may differ in their sleep-dependent consolidation. Three options come to mind. First, initial differences in memory strength could drive subsequent differences in sleep-related consolidation. In our paradigm, initial learning favored a ‘one-shot’ approach to model a natural memory acquisition process, without extensive rehearsal. With this approach, after learning, the initial memory strength of recognition memory (∼89%) was higher than that of cued association (∼38%). Since a new memory must pass a certain minimal level of performance to be able to further benefit from sleep (Drosopoulos et al., 2007; Schoch et al., 2017), these initial differences in memory strength could have driven subsequent sleep-dependent consolidation differences. This aspect might have been magnified by the task design. Participants had to repeat the pet name during encoding and thus focused on the pet rather than on the association itself. In addition, our paradigm included an immediate retrieval task (not always employed in sleep and memory studies), which constitutes a re-consolidation step involving intentional re-activations (‘testing effect’, (Rowland, 2014; Sutterer & Awh, 2016). Intentional rehearsal would likely mostly affect recognition because of the high retention rate at the immediate retrieval. A second option is that a short nap is not sufficient to significantly strengthen the relational aspect of memory association, which may require longer consolidation than that needed for recognition memory. However, sleep effects in the cued associate retrieval were also not detectable in the vPAL paradigm after longer sleep periods in epilepsy patients (Geva-Sagiv et al., 2023). Third, sleep may preferentially stabilize aspects of memory connected to prior knowledge. Evidence from rodents (Tse et al., 2007) and humans (Hennies et al., 2016; Kesteren et al., 2013) suggest that prior knowledge to which learning items can be connected facilitates memory consolidation. This has even been demonstrated for targeted memory reactivation (Groch et al., 2017). In the vPAL task, we used pictures of easily recognizable celebrities that could have biased the consolidation towards these images, relative to the newly learned association between celebrities and pets. Alternatively, competing models for memory stabilization such as contextual binding theory could explain these results, if sleep reduced contextual interference (and thus improving recognition), rather than strengthening the associations (Yonelinas et al., 2019). Additional studies are needed to determine whether and how memory accuracy and association differ in their sleep-dependent consolidation profiles.

Future studies with vPAL could expand this investigation along several avenues. Adding another memory task may help clarify further what aspects of declarative memory benefit from sleep. For example, a relational memory task could help focus on the associative aspects of the memory which may particularly depend on hippocampal activity (Bunsey & Eichenbaum, 1996). In addition, this task could be further combined with neuroimaging or intracranial monitoring of brain activity to further determine the neural processes mediating sleep-dependent memory consolidation. Finally, future research will determine the utility of vPAL in other populations such as the elderly and patients with amnesia, epilepsy, or cognitive impairment due to neurodegeneration, and, with time, may be used to investigate methods for boosting memory, particularly in people with learning and memory impairments.

## Methods

### Participants

Overall, 32 participants (average age = 27.0 years, range: 22 to 37 years, 39.3% female) were each tested on two separate experimental days. All participants were between 22 and 37 years old, without a history or presence of neuropsychiatric or sleep disorders. Written informed consent was obtained from each participant. The study was approved by the Medical Institutional Review Board at the Tel Aviv Sourasky Medical Center. Two participants were excluded because the EEG revealed that they could not sleep at all in the ‘sleep’ session, and one participant was excluded due to insufficient quality of the electrophysiology recording. Furthermore, one participant was excluded due to general memory issues (chance performance in all memory tasks), resulting in 28 participants available for recognition memory measures. Due to technical issues, the cued recall (audio recordings of verbal reports) of 5 participants was corrupted, resulting in 23 participants available for association memory measures.

### Experimental procedure

Each participant participated in two visual paired associative learning (vPAL) experimental sessions, conducted one week apart at the same time of day (around noon), each consisting of an encoding (learning) phase and two retrieval tests (**Figure 1**). We ran a 2x2 within-subject design (2 time points x 2 session types) such that memory performance was assessed 4 times in each participant (without any feedback). The two retrieval tests were separated by 2.5 hours when subjects were either given an opportunity to sleep (daytime nap with EEG, details below) or rest awake in the lab, thereby forming either a learning-test-wake-test or a learning-test-sleep-test schedule. The order of sleep and wakefulness sessions was counterbalanced across participants (either wake first or sleep first), and different image sets were used for each session (their assignment to either the first or second sessions was also counterbalanced). Participants were instructed to avoid drinking coffee and sleep in the 6h before each session. Each session consisted of the following steps: (a) learning: encoding of image-pair associations, (b) immediate retrieval: recognition and cued recall of learned associations, (c) a consolidation period of 2.5h between the two retrieval task when participants either stayed awake or were given an opportunity to sleep, and (d) delayed retrieval: identical to the immediate retrieval step testing recognition and association recall. The constant schedule in each session helps focus on the comparison between sleep and wake conditions and go beyond the effects of learning or circadian effects of time of day, which are similarly present in both sleep and wake sessions.

#### vPAL encoding phase

Participants were asked to memorize a series of 25 image pairs. Each pair consisted of a celebrity (e.g. Amy Winehouse) and their pet (e.g. dog) and participants were asked to learn which pet belongs to which celebrity (“Try to memorize the pet for each celebrity”). Each image pair was presented for 2s. To ensure that subjects are attentive, that the association is encoded, and to ensure that the pets are recognized correctly, participants repeated the pet’s name out loud. The learning phase enabled the encoding of 25 associations within 15min. Images of celebrities were used for quick, one-trial learning because they are easy to remember (memorability of emotional vs neutral memory, LaBar & Cabeza, 2006; Wagner et al., 2001). The downfall of having a quick encoding of only 25 associations is a high subject-variability, which we addressed by a within-subject design (two sessions per subject).

#### vPAL retrieval phase

Participants underwent two identical retrieval tests, before (‘immediate retrieval’) and after (‘delayed retrieval’) the consolidation period. During the retrieval task, single images of celebrities were presented for two seconds and subsequently queried (40 images in total). Of these images, 25 celebrities were images that were originally presented and paired with a pet during the learning phase and thus represent pet-owners. 15 additional images of celebrities were new (‘lures’). The same set of lures was used in the immediate and the delayed retrieval. We separately probed two different aspects of memory retrieval: (i) the accuracy of memory recognition where participants were asked to identify pet-owners and reject lures (“Is this a pet owner?”, Yes/No) and, only if a celebrity was identified as a pet-owner, (ii) participants then concluded a cued recall by naming the associated pet (“name the pet”) while their verbal response was registered. The cued recall (ii-retrieval) was skipped if a participant did not recognize the celebrity. Overall, we separately estimated which of the 40 queried images were correctly identified as new or old and whether the participants were able to recollect the associated pet (when applicable). Importantly, feedback was never provided, neither during the retrieval task before nor after the consolidation period, and thus the first retrieval task before the consolidation did not function as explicit re-learning.

To determine how memory consolidation in the vPAL paradigm is affected by sleep, we measured memory retrieval scores four times: before and after a 2.5h period when subjects were either awake or given an opportunity to sleep (Figure 2A,B). To further quantify memory effects, we computed memory consolidation (by immediate and delayed retrieval tests), and further compared consolidation scores between sleep and wake rest consolidation (Figure 2C,D). To this end, we defined two measures of memory consolidation as follows:

i. Recognition discrimination = z(# correct recognition / N_trials_) - z(# false recognition /N_trials_); in signal detection theory referred to as D-prime (z= z-transform, Macmillan & Creelman, 1991). Then, memory stability score (Δ Recognition discrimination stability) was computed per session as the difference in % between delayed and immediate retrieval tests: recognition memory _delayed_ - recognition memory _immediate_. This stability score was compared between sleep and wake sessions. Thus, the difference between the Δ Recognition discrimination stability in sleep and Δ Recognition discrimination stability in wake represents the added benefit of sleep for consolidation of recognition discrimination, reflecting stabilizing effects of sleep on memory (Rasch & Born, 2013; Schreiner & Staudigl, 2020; Walker & Stickgold, 2004)
ii. Correct associations = # correct cued recall answers / N _total_trials_. Then association stability score (Δ association stability) was computed per session as the difference in % between delayed and immediate retrieval tests: association accuracy _delayed_ - association accuracy _immediate_. This stability score was compared between sleep and wake sessions. Thus, the difference between the Δ association stability in sleep and Δ association stability in wake represents the added benefit of sleep for retrieving associations in a cued recall. N _total_trials_ was always set to 25, including not recognized celebrities, no answers, and incorrectly paired pets.

### Images and visual stimulation

Images of celebrities and animals were generated from copyright-free web images. We created 2 image sets for vPAL experiments, each with 25 image pairs of celebrities and animals for learning, and 15 additional pictures of lure celebrities. Different sets were used during sleep and wake sessions such that no celebrity or animal appeared twice. The set’s allocation to a session was counterbalanced. Hence, each image was presented an equal number of times, though in different participants. Altogether, this represents a total of 130 images used in this study (80 celebrities and 50 animals). Because of the within-subject within-session comparisons, there was no need for shuffling the learned-lure image allocation. But we randomized the order of the stimuli presentation to control for order effects. Stimuli were presented using the psychtoolbox implementation in Python (http://psychtoolbox) on a monitor placed about 50 cm in front of participants.

### Sleep EEG, sleep scoring, and sleep spindle detection

#### Data collection

In sleep sessions, we collected polysomnographic data including high-density electroencephalogram (hd-EEG), electrooculogram (EOG), electromyogram (EMG), electrocardiogram, and video. hd-EEG was recorded using a 256-channel hydrogel geodesic sensor net (Electrical Geodesics, Inc. [EGI]). Signals were referred to Cz, amplified using an antialiasing filter and an AC-coupled high input impedance amplifier (NetAmps 300, EGI), and then digitized at 1000 Hz. Before the recording began, the electrode impedance of all sensors was confirmed to be at least below 50 kΩ.

#### Sleep scoring and analysis

Sleep scoring was performed according to the established guidelines of the American Academy of Sleep Medicine (Iber et al., 2007) with the aid of the sleep module in the Visbrain Python package (Combrisson et al., 2019). To this end, EEG data from F3/F4, C3/C4, and O1/O2 was re-referenced to the contralateral mastoid, resampled to 250 Hz, band-pass filtered to 0.3-40 Hz and visualized in 30s epochs together with synchronized EOG from electrodes E1&E2 above and below eyebrows and EMG from submental electrode (band-pass filtered to 10-100 Hz). Successful scoring was further verified by inspecting the time-frequency representation (spectrogram) of the Pz electrode (not involved in the scoring process) superimposed with the hypnogram. Sleep was characterized, for each participant, by extracting the following measures: time in bed, duration from start to end of sleep, duration of wake within sleep, total sleep time, time in each sleep state, time in NREM sleep, latency to onset of each sleep stage, sleep onset latency, sleep efficiency (time asleep / time in bed) and sleep maintenance (time asleep / start to end sleep).

#### Sleep spindle detection

We detected sleep spindles automatically with the Python-based YASA toolbox (Vallat & Walker, 2021). The EEG signal was cleaned and preprocessed as follows: we resampled the signal to a sample rate of 250 Hz and bandpass-filtered it in the range of 0.3 to 40 Hz. We then interpolated noisy channels (6.06 ± 2.51%) replacing the signal with an average of adjacent electrodes and excluded bad epochs with artifacts (6.14 ± 3.36%) from further analysis via manual labeling. We then re-referenced the signal to the global average and visually confirmed the data’s quality in each participant by looking at the PSD and the scalp topography of sleep spindle activity. We detected potential spindles events when the signal exceeded the thresholds for the YASA criteria of relative power (ratio > 0.2), signal to sigma-envelope correlation (r > 0.65), root mean square of sigma-envelope (rms > mean + 1.5 SD). Furthermore, we excluded events within 500ms of other events, or when duration was not in the 0.5-2s range. We performed spindle detection separately in 15 electrodes comprising a centro-parietal region of interest (Figure 3E) and computed the participant’s spindle density as the average of observed densities in those electrodes. Spindle density was defined as the number of detected spindles in N2 and N3 divided by the minutes spent in N2 and N3. Finally, we computed the Pearson correlation between individual average spindle density and the memory decay in the recognition task (during sleep). Due to a corrupted file, we had to drop one more participant for the spindle detection.

### Statistical analysis

We assessed the normality of distributions with a Shapiro-Wilk test. Parametric methods for statistical testing were used if the data were normally distributed. For the analysis of the recognition answers, we used Wilcoxon signed-rank test, because the data was not normally distributed. We applied two-sided tests unless stated otherwise. Data processing and analysis were performed using custom-made R® scripts (R Core Team, 2021).

## Acknowledgments

We thank Gal Zatelman, May Eliyahu, Tomer Cohen, and Romario Zarik for assistance with data collection. Dr. Noa Bar-Ilan Regev for administrative assistance. Charan Ranganath, Fried and Nir lab members for discussions and input. Avi Mendelsohn and Hagar Gelbard-Sagiv for advice on paradigm development. Supported by US National Science Foundation and U.S.-Israel Binational Science Foundation NSF-BSF grant 2017628 (I.F. & Y.N.), ERC-2019-CoG 864353 (Y.N.)., M.G.S. was funded by a Postdoctoral fellowship from the Rothschild Foundation and a Tel Aviv University Sagol School of Neuroscience Postdoctoral Fellowship.

## Author contributions

Y.N., M.G.S, and I.F. conceived the research and designed experiments. Y.N. and I.F. secured funding. M.G.S., R.F., and O.S. prepared the cognitive experiment. R.F., and Y.G. managed data collection. Y.G., R.F., S.Y., and F.J.S. performed sleep scoring. F.J.S., R.F., and S.Y. performed data analysis. F.J.S. and Y.N. wrote the manuscript. All authors provided ongoing critical review of results and commented on the manuscript.

## Data availability

The data underlying this article will be shared on reasonable request to the corresponding author.

## References

Andrillon, T., Nir, Y., Staba, R. J., Ferrarelli, F., Cirelli, C., Tononi, G., & Fried, I. (2011). Sleep spindles in humans: Insights from intracranial EEG and unit recordings. The Journal of Neuroscience: The Official Journal of the Society for Neuroscience, 31(49), 17821–17834. 10.1523/JNEUROSCI.2604-11.2011

Antony, J. W., Schönauer, M., Staresina, B. P., & Cairney, S. A. (2019). Sleep Spindles and Memory Reprocessing. Trends in Neurosciences, 42(1), 1–3. 10.1016/j.tins.2018.09.012

Backhaus, J., & Junghanns, K. (2006). Daytime naps improve procedural motor memory. Sleep Medicine, 7(6), 508–512. 10.1016/j.sleep.2006.04.002

Bailes, C., Caldwell, M., Wamsley, E. J., & Tucker, M. A. (2020). Does sleep protect memories against interference? A failure to replicate. PLOS ONE, 15(2), e0220419. 10.1371/journal.pone.0220419

Bar, E., Marmelshtein, A., Arzi, A., Perl, O., Livne, E., Hizmi, E., Paz, R., Sobel, N., Dudai, Y., & Nir, Y. (2020). Local Targeted Memory Reactivation in Human Sleep. Current Biology, 30(8), 1435–1446.e5. 10.1016/j.cub.2020.01.091

Barrett, T. R., & Ekstrand, B. R. (1972). Effect of sleep on memory: III. Controlling for time-of-day effects. Journal of Experimental Psychology, 96, 321–327. 10.1037/h0033625

Bergmann, T. O., Mölle, M., Diedrichs, J., Born, J., & Siebner, H. R. (2012). Sleep spindle-related reactivation of category-specific cortical regions after learning face-scene associations. NeuroImage, 59(3), 2733–2742. 10.1016/j.neuroimage.2011.10.036

Bird, C. M. (2017). The role of the hippocampus in recognition memory. Cortex, 93, 155–165. 10.1016/j.cortex.2017.05.016

Bunsey, M., & Eichenbaum, H. (1996). Conservation of hippocampal memory function in rats and humans. Nature, 379(6562), Article 6562. 10.1038/379255a0

Buzsáki, G. (1989). Two-stage model of memory trace formation: A role for “noisy” brain states. Neuroscience, 31(3), 551–570. 10.1016/0306-4522(89)90423-5

Cairney, S. A., Guttesen, A. á V., El Marj, N., & Staresina, B. P. (2018). Memory Consolidation Is Linked to Spindle-Mediated Information Processing during Sleep. Current Biology, 28(6), 948–954.e4. 10.1016/j.cub.2018.01.087

Combrisson, E., Vallat, R., O’Reilly, C., Jas, M., Pascarella, A., Saive, A., Thiery, T., Meunier, D., Altukhov, D., Lajnef, T., Ruby, P., Guillot, A., & Jerbi, K. (2019). Visbrain: A Multi-Purpose GPU-Accelerated Open-Source Suite for Multimodal Brain Data Visualization. Frontiers in Neuroinformatics, 13. https://www.frontiersin.org/articles/10.3389/fninf.2019.00014

Cordi, M. J., & Rasch, B. (2021). How robust are sleep-mediated memory benefits? Current Opinion in Neurobiology, 67, 1–7. 10.1016/j.conb.2020.06.002

Datta, S., & MacLean, R. R. (2007). Neurobiological mechanisms for the regulation of mammalian sleep– wake behavior: Reinterpretation of historical evidence and inclusion of contemporary cellular and molecular evidence. Neuroscience & Biobehavioral Reviews, 31(5), 775–824. 10.1016/j.neubiorev.2007.02.004

Diekelmann, S., & Born, J. (2010). The memory function of sleep. Nature Reviews Neuroscience, 11(2), Article 2. 10.1038/nrn2762

Drosopoulos, S., Schulze, C., Fischer, S., & Born, J. (2007). Sleep’s function in the spontaneous recovery and consolidation of memories. Journal of Experimental Psychology: General, 136, 169–183. 10.1037/0096-3445.136.2.169

Dudai, Y., Karni, A., & Born, J. (2015). The Consolidation and Transformation of Memory. Neuron, 88(1), 20–32. 10.1016/j.neuron.2015.09.004

Ebbinghaus, H. (1913). Memory: A contribution to experimental psychology. *Eachers College*, Columbia University, 20(4), 155–156. 10.5214/ans.0972.7531.200408

Ellenbogen, J. M., Hu, P. T., Payne, J. D., Titone, D., & Walker, M. P. (2007). Human relational memory requires time and sleep. Proceedings of the National Academy of Sciences, 104(18), 7723–7728. 10.1073/pnas.0700094104

Ellenbogen, J. M., Hulbert, J. C., Stickgold, R., Dinges, D. F., & Thompson-Schill, S. L. (2006). Interfering with Theories of Sleep and Memory: Sleep, Declarative Memory, and Associative Interference. Current Biology, 16(13), 1290–1294. 10.1016/j.cub.2006.05.024

Fischer, S., Hallschmid, M., Elsner, A. L., & Born, J. (2002). Sleep forms memory for finger skills. Proceedings of the National Academy of Sciences, 99(18), 11987–11991. 10.1073/pnas.182178199

Frankland, P. W., & Bontempi, B. (2005). The organization of recent and remote memories. Nature Reviews Neuroscience, 6(2), Article 2. 10.1038/nrn1607

Gais, S., Lucas, B., & Born, J. (2006). Sleep after learning aids memory recall. Learning & Memory, 13(3), 259–262. 10.1101/lm.132106

Gais, S., Mölle, M., Helms, K., & Born, J. (2002). Learning-Dependent Increases in Sleep Spindle Density. Journal of Neuroscience, 22(15), 6830–6834. 10.1523/JNEUROSCI.22-15-06830.2002

Geva-Sagiv, M., Mankin, E., Eliashiv, D., Epstein, S., Cherry, N., Kalender, G., Tchemodanov, N., Nir, Y., & Fried, I. (2023). Augmenting hippocampal-prefrontal neuronal synchrony in human sleep enhances memory consolidation. Nature Neuroscience.

Groch, S., Schreiner, T., Rasch, B., Huber, R., & Wilhelm, I. (2017). Prior knowledge is essential for the beneficial effect of targeted memory reactivation during sleep. Scientific Reports, 7, 39763. 10.1038/srep39763

Hall, J. F. (1971). Verbal learning, retention, and memory. Canadian Journal of Psychology/Revue Canadienne de Psychologie, 25(5), 412.

Hennies, N., Lambon Ralph, M. A., Kempkes, M., Cousins, J. N., & Lewis, P. A. (2016). Sleep Spindle Density Predicts the Effect of Prior Knowledge on Memory Consolidation. The Journal of Neuroscience, 36(13), 3799–3810. 10.1523/JNEUROSCI.3162-15.2016

Iber, C., Ancoli-Israel, S., Chesson, A., & Quan, S. F. (2007). The AASM Manual for the Scoring of Sleep and Associated Events: Rules, Terminology and Technical Specifications. American Academy of Sleep Medicine.

Jenkins, J. G., & Dallenbach, K. M. (1924). Obliviscence during Sleep and Waking. The American Journal of Psychology, 35(4), 605–612. 10.2307/1414040

Káli, S., & Dayan, P. (2004). Off-line replay maintains declarative memories in a model of hippocampal-neocortical interactions. Nature Neuroscience, 7(3), Article 3. 10.1038/nn1202

Karni, A., Tanne, D., Rubenstein, B. S., Askenasy, J. J. M., & Sagi, D. (1994). Dependence on REM Sleep of Overnight Improvement of a Perceptual Skill. Science, 265(5172), 679–682. 10.1126/science.8036518

Kesteren, M. T. R. van, Rijpkema, M., Ruiter, D. J., & Fernández, G. (2013). Consolidation Differentially Modulates Schema Effects on Memory for Items and Associations. PLOS ONE, 8(2), e56155. 10.1371/journal.pone.0056155

Korman, M., Doyon, J., Doljansky, J., Carrier, J., Dagan, Y., & Karni, A. (2007). Daytime sleep condenses the time course of motor memory consolidation. Nature Neuroscience, 10(9), Article 9. 10.1038/nn1959

LaBar, K. S., & Cabeza, R. (2006). Cognitive neuroscience of emotional memory. Nature Reviews Neuroscience, 7(1), Article 1. 10.1038/nrn1825

Lahl, O., Wispel, C., Willigens, B., & Pietrowsky, R. (2008). An ultra short episode of sleep is sufficient to promote declarative memory performance. Journal of Sleep Research, 17(1), 3–10. 10.1111/j.1365-2869.2008.00622.x

Lau, H., Tucker, M. A., & Fishbein, W. (2010). Daytime napping: Effects on human direct associative and relational memory. Neurobiology of Learning and Memory, 93(4), 554–560. 10.1016/j.nlm.2010.02.003

Macmillan, N. A., & Creelman, C. D. (1991). Detection theory: A user’s guide (pp. xv, 407). Cambridge University Press.

Maquet, P. (2001). The Role of Sleep in Learning and Memory. Science, 294(5544), 1048–1052. 10.1126/science.1062856

Marshall, L., & Born, J. (2007). The contribution of sleep to hippocampus-dependent memory consolidation. Trends in Cognitive Sciences, 11(10), 442–450. 10.1016/j.tics.2007.09.001

McClelland, J. L., McNaughton, B. L., & O’Reilly, R. C. (1995). Why there are complementary learning systems in the hippocampus and neocortex: Insights from the successes and failures of connectionist models of learning and memory. Psychological Review, 102, 419–457. 10.1037/0033-295X.102.3.419

McDermott, K. B., Szpunar, K. K., & Christ, S. E. (2009). Laboratory-based and autobiographical retrieval tasks differ substantially in their neural substrates. Neuropsychologia, 47(11), 2290–2298. 10.1016/j.neuropsychologia.2008.12.025

Mednick, S. C., McDevitt, E. A., Walsh, J. K., Wamsley, E., Paulus, M., Kanady, J. C., & Drummond, S. P. A. (2013). The Critical Role of Sleep Spindles in Hippocampal-Dependent Memory: A Pharmacology Study. Journal of Neuroscience, 33(10), 4494–4504. 10.1523/JNEUROSCI.3127-12.2013

Mednick, S., Nakayama, K., & Stickgold, R. (2003). Sleep-dependent learning: A nap is as good as a night. Nature Neuroscience, 6(7), Article 7. 10.1038/nn1078

Muehlroth, B. E., Sander, M. C., Fandakova, Y., Grandy, T. H., Rasch, B., Shing, Y. L., & Werkle-Bergner, M. (2019). Precise Slow Oscillation–Spindle Coupling Promotes Memory Consolidation in Younger and Older Adults. Scientific Reports, 9(1), Article 1. 10.1038/s41598-018-36557-z

Nádasdy, Z., Hirase, H., Czurkó, A., Csicsvari, J., & Buzsáki, G. (1999). Replay and Time Compression of Recurring Spike Sequences in the Hippocampus. Journal of Neuroscience, 19(21), 9497–9507. 10.1523/JNEUROSCI.19-21-09497.1999

Newbury, C. R., & Monaghan, P. (2019). When does sleep affect veridical and false memory consolidation? A meta-analysis. Psychonomic Bulletin & Review, 26(2), 387–400. 10.3758/s13423-018-1528-4

Niknazar, H., Malerba, P., & Mednick, S. C. (2022). Slow oscillations promote long range effective communication: The key for memory consolidation in a broken down network (p. 2022.04.13.488133). bioRxiv. 10.1101/2022.04.13.488133

Nir, Y., Staba, R. J., Andrillon, T., Vyazovskiy, V. V., Cirelli, C., Fried, I., & Tononi, G. (2011). Regional slow waves and spindles in human sleep. Neuron, 70(1), 153–169. 10.1016/j.neuron.2011.02.043

Nishida, M., Pearsall, J., Buckner, R. L., & Walker, M. P. (2009). REM Sleep, Prefrontal Theta, and the Consolidation of Human Emotional Memory. Cerebral Cortex, 19(5), 1158–1166. 10.1093/cercor/bhn155

Nishida, M., & Walker, M. P. (2007). Daytime Naps, Motor Memory Consolidation and Regionally Specific Sleep Spindles. PLOS ONE, 2(4), e341. 10.1371/journal.pone.0000341

Payne, J. D., Stickgold, R., Swanberg, K., & Kensinger, E. A. (2008). Sleep Preferentially Enhances Memory for Emotional Components of Scenes. Psychological Science, 19(8), 781–788. 10.1111/j.1467-9280.2008.02157.x

Petzka, M., Chatburn, A., Charest, I., Balanos, G. M., & Staresina, B. P. (2022). Sleep spindles track cortical learning patterns for memory consolidation. Current Biology, 32(11), 2349–2356.e4. 10.1016/j.cub.2022.04.045

Peyrache, A., Khamassi, M., Benchenane, K., Wiener, S. I., & Battaglia, F. P. (2009). Replay of rule-learning related neural patterns in the prefrontal cortex during sleep. Nature Neuroscience, 12(7), Article 7. 10.1038/nn.2337

Plihal, W., & Born, J. (1997). Effects of Early and Late Nocturnal Sleep on Declarative and Procedural Memory. Journal of Cognitive Neuroscience, 9(4), 534–547. 10.1162/jocn.1997.9.4.534

Pöhlchen, D., Pawlizki, A., Gais, S., & Schönauer, M. (2021). Evidence against a large effect of sleep in protecting verbal memories from interference. Journal of Sleep Research, 30(2), e13042. 10.1111/jsr.13042

Quiroga, R. Q., Reddy, L., Kreiman, G., Koch, C., & Fried, I. (2005). Invariant visual representation by single neurons in the human brain. Nature, 435(7045), Article 7045. 10.1038/nature03687

R Core Team. (2021). R: A Language and Environment for Statistical Computing [Computer software]. R Foundation for Statistical Computing. https://www.R-project.org/

Rasch, B., & Born, J. (2013). About Sleep’s Role in Memory. Physiological Reviews, 93(2), Article 2. 10.1152/physrev.00032.2012

Rasch, B., Büchel, C., Gais, S., & Born, J. (2007). Odor Cues During Slow-Wave Sleep Prompt Declarative Memory Consolidation. Science, 315(5817), 1426–1429. 10.1126/science.1138581

Robertson, E. M., Pascual-Leone, A., & Miall, R. C. (2004). Current concepts in procedural consolidation. Nature Reviews Neuroscience, 5(7), Article 7. 10.1038/nrn1426

Rowland, C. A. (2014). The effect of testing versus restudy on retention: A meta-analytic review of the testing effect. Psychological Bulletin, 140, 1432–1463. 10.1037/a0037559

Ruch, S., Markes, O., Duss, S. B., Oppliger, D., Reber, T. P., Koenig, T., Mathis, J., Roth, C., & Henke, K. (2012). Sleep stage II contributes to the consolidation of declarative memories. Neuropsychologia, 50(10), Article 10. 10.1016/j.neuropsychologia.2012.06.008

Rudoy, J. D., Voss, J. L., Westerberg, C. E., & Paller, K. A. (2009). Strengthening Individual Memories by Reactivating Them During Sleep. Science, 326(5956), 1079–1079. 10.1126/science.1179013

Schabus, M., Gruber, G., Parapatics, S., Sauter, C., Klösch, G., Anderer, P., Klimesch, W., Saletu, B., & Zeitlhofer, J. (2004). Sleep spindles and their significance for declarative memory consolidation. Sleep, 27(8), 1479–1485. 10.1093/sleep/27.7.1479

Schoch, S. F., Cordi, M. J., & Rasch, B. (2017). Modulating influences of memory strength and sensitivity of the retrieval test on the detectability of the sleep consolidation effect. Neurobiology of Learning and Memory, 145, 181–189. 10.1016/j.nlm.2017.10.009

Schönauer, M., Alizadeh, S., Jamalabadi, H., Abraham, A., Pawlizki, A., & Gais, S. (2017). Decoding material-specific memory reprocessing during sleep in humans. Nature Communications, 8, 15404. 10.1038/ncomms15404

Schreiner, T., & Staudigl, T. (2020). Electrophysiological signatures of memory reactivation in humans. Philosophical Transactions of the Royal Society B: Biological Sciences, 375(1799), 20190293. 10.1098/rstb.2019.0293

Seibt, J., Richard, C. J., Sigl-Glöckner, J., Takahashi, N., Kaplan, D. I., Doron, G., de Limoges, D., Bocklisch, C., & Larkum, M. E. (2017). Cortical dendritic activity correlates with spindle-rich oscillations during sleep in rodents. Nature Communications, 8(1), Article 1. 10.1038/s41467-017-00735-w

Sharon, O., & Nir, Y. (2018). Attenuated Fast Steady-State Visual Evoked Potentials During Human Sleep. *Cerebral Cortex (New York*, N.Y.: 1991), *28*(4), 1297–1311. 10.1093/cercor/bhx043

Smith, C. N., Jeneson, A., Frascino, J. C., Kirwan, C. B., Hopkins, R. O., & Squire, L. R. (2014). When recognition memory is independent of hippocampal function. Proceedings of the National Academy of Sciences, 111(27), 9935–9940. 10.1073/pnas.1409878111

Steriade, M., & Timofeev, I. (2003). Neuronal Plasticity in Thalamocortical Networks during Sleep and Waking Oscillations. Neuron, 37(4), 563–576. 10.1016/S0896-6273(03)00065-5

Studte, S., Bridger, E., & Mecklinger, A. (2015). Nap sleep preserves associative but not item memory performance. Neurobiology of Learning and Memory, 120, 84–93. 10.1016/j.nlm.2015.02.012

Suthana, N. A., Parikshak, N. N., Ekstrom, A. D., Ison, M. J., Knowlton, B. J., Bookheimer, S. Y., & Fried, I. (2015). Specific responses of human hippocampal neurons are associated with better memory. Proceedings of the National Academy of Sciences, 112(33), 10503–10508. 10.1073/pnas.1423036112

Sutterer, D. W., & Awh, E. (2016). Retrieval practice enhances the accessibility but not the quality of memory. Psychonomic Bulletin & Review, 23(3), 831–841. 10.3758/s13423-015-0937-x

Tononi, G., & Cirelli, C. (2014). Sleep and the Price of Plasticity: From Synaptic and Cellular Homeostasis to Memory Consolidation and Integration. Neuron, 81(1), 12–34. 10.1016/j.neuron.2013.12.025

Tse, D., Langston, R. F., Kakeyama, M., Bethus, I., Spooner, P. A., Wood, E. R., Witter, M. P., & Morris, R. G. M. (2007). Schemas and memory consolidation. *Science (New York*, N.Y.), 316(5821), 76–82. 10.1126/science.1135935

Tucker, M. A., Hirota, Y., Wamsley, E. J., Lau, H., Chaklader, A., & Fishbein, W. (2006). A daytime nap containing solely non-REM sleep enhances declarative but not procedural memory. Neurobiology of Learning and Memory, 86(2), 241–247. 10.1016/j.nlm.2006.03.005

Vallat, R., & Walker, M. P. (2021). An open-source, high-performance tool for automated sleep staging. eLife, 10, e70092. 10.7554/eLife.70092

Wagner, U., Gais, S., & Born, J. (2001). Emotional Memory Formation Is Enhanced across Sleep Intervals with High Amounts of Rapid Eye Movement Sleep. Learning & Memory, 8(2), 112–119. 10.1101/lm.36801

Walker, M. P., & Stickgold, R. (2004). Sleep-Dependent Learning and Memory Consolidation. Neuron, 44(1), 121–133. 10.1016/j.neuron.2004.08.031

Walker, M. P., Stickgold, R., Alsop, D., Gaab, N., & Schlaug, G. (2005). Sleep-dependent motor memory plasticity in the human brain. Neuroscience, 133(4), 911–917. 10.1016/j.neuroscience.2005.04.007

Werchan, D. M., & Gómez, R. L. (2013). Generalizing memories over time: Sleep and reinforcement facilitate transitive inference. Neurobiology of Learning and Memory, 100, 70–76. 10.1016/j.nlm.2012.12.006

Yonelinas, A. P., Ranganath, C., Ekstrom, A. D., & Wiltgen, B. J. (2019). A contextual binding theory of episodic memory: Systems consolidation reconsidered. Nature Reviews Neuroscience, 20(6), 364–375. 10.1038/s41583-019-0150-4

Zeitlhofer, J., Gruber, G., Anderer, P., Asenbaum, S., Schimicek, P., & Saletu, B. (1997). Topographic distribution of sleep spindles in young healthy subjects. Journal of Sleep Research, 6(3), 149–155. 10.1046/j.1365-2869.1997.00046.x

